# Degenerative and regenerative peripheral processes are associated with persistent painful chemotherapy-induced neuropathies in males and females

**DOI:** 10.1101/2024.01.25.577218

**Authors:** George T. Naratadam, Jennifer Mecklenburg, Sergey A. Shein, Yi Zou, Zhao Lai, Alexei V. Tumanov, Theodor J. Price, Armen N. Akopian

## Abstract

This study aimed to investigate the time course of gene expression changes during the progression of persistent painful neuropathy caused by paclitaxel (PTX) in male and female mouse hind paws and dorsal root ganglia (DRG). Bulk RNA-seq was used to investigate the gene expression changes in the paw and DRG collected at 1, 16, and 31 days post-PTX. At these time points, differentially expressed DEGs were predominantly related to reduction or increase in epithelial, skin, bone, and muscle development and to angiogenesis, myelination, axonogenesis, and neurogenesis. These processes were accompanied by regulation of DEGs related to cytoskeleton, extracellular matrix organization and cellular energy production. This gene plasticity during persistent painful neuropathy progression likely represents biological processes linked to tissue regeneration and degeneration. Unlike regeneration/degeneration, gene plasticity related to immune processes was minimal at 1–31 days post-PTX. It was also noted that despite similarities in biological processes and pain chronicity in males and females, specific DEGs showed dramatic sex-dependency. The main conclusions of this study are that gene expression plasticity in paws and DRG during PTX neuropathy progression relates to tissue regeneration and degeneration, minimally affects the immune system processes, and is heavily sex-dependent at the individual gene level.

## Introduction

Chronic pain conditions can develop through several phases: initiation of pain, pain maintenance, and finally, either resolution or persistency of hypersensitivity/pain. Bulk RNA-sequencing (RNA-seq) experiments show that pain conditions trigger substantial gene expression plasticity in nociceptive pathway tissues, including the DRG and spinal cord(Cao et al., 2019; North et al., 2019; Tang et al., 2020; Guo et al., 2021; Li et al., 2021a; Li et al., 2021b; Doty et al., 2022; Xu et al., 2022). Despite this progress, there are still several gaps in knowledge. First, gene expression plasticity in peripheral tissues, such as the paw and muscles, during pain conditions has not been well addressed. Second, gene plasticity is probably chronic pain condition phase-dependent, and this area is grossly understudied. Third, the higher prevalence of pain conditions in women suggests that there are likely mechanistic sex difference in chronic pain that are also not well understood (Unruh, 1996; Berkley, 1997; Fillingim et al., 2009; Traub and Ji, 2013; Ray et al., 2023). While there is a consensus that gene expression plasticity in the peripheral and central portions of nociceptive pathways can be sex-dependent(Mecklenburg et al., 2020), potential sex differences in gene expression occurring at different phases of neuropathic pain development and maintenance is understudied.

Based on the above outlined critical gaps in knowledge, the aims of this study were to investigate the time course of gene expression plasticity in male and female mouse hind paws and DRG during the progression of persistent painful neuropathy. The rationale for these aims was to understand what types of gene expression plasticity are associated with different phases of persistent painful neuropathies; and whether these changes were sex-dependent. We used a paclitaxel (PTX)-induced peripheral neuropathy model to produce persistent pain in both male and female mice. Transcriptomic profiling was used to measure gene expression changes at different post-PTX time points to dissect the biological processes accompanying the development of pain persistency in males and females.

## Materials and Methods

### Ethical Approval

The reporting in the manuscript follows the recommendations in the ARRIVE guidelines (PLoS Bio 8(6), e1000412,e1000412,2010). We also followed guidelines issued by the National Institutes of Health (NIH) and the Society for Neuroscience (SfN) to minimize the number of animals used and their suffering. All animal experiments conformed to protocols approved by the University Texas Health Science Center at San Antonio (UTHSCSA) Institutional Animal Care and Use Committee (IACUC). Protocol numbers are 20180001AR and 20200011AR.

### Reagents and mouse lines

Paclitaxel (PTX) was purchased from Millipore-Sigma (Cat: T7402; St. Louis, MO). PTX powder was replaced every 4 months to maintain the reagent’s activity at the same level. PTX stock (40mg/ml) was prepared in 100% ethanol, which was kept in -80°C for no more than 2 weeks. Stock was diluted to working concentrations using a mixture of 650 ml saline/160 ml Kolliphor-EL (Millipore-Sigma; Cat: C5135; St. Louis, MO). Remnant diluted stock was discarded after every injection.

Experiments were performed on 10–18-week-old C57BL/6 wild-type male and female mice, which were originally purchased from Jackson Laboratory (Bar Harbor, ME).

### Paclitaxel (PTX)-induced pain model and measurement of hypersensitivity

PTX was injected systemically (i.p.). An appropriate sample size for each group was computed using power analysis (see below). PTX was titrated for achieving persistent (>50 days) painful chemotherapy-induced models. PTX was injected three times at -6, -3, and 0 days at 6 mg/kg dosage to generate persistent mechanical hypersensitivity for at least 50 days (see schematic; *Figs 1A, 1B*).

**Figure 1:**
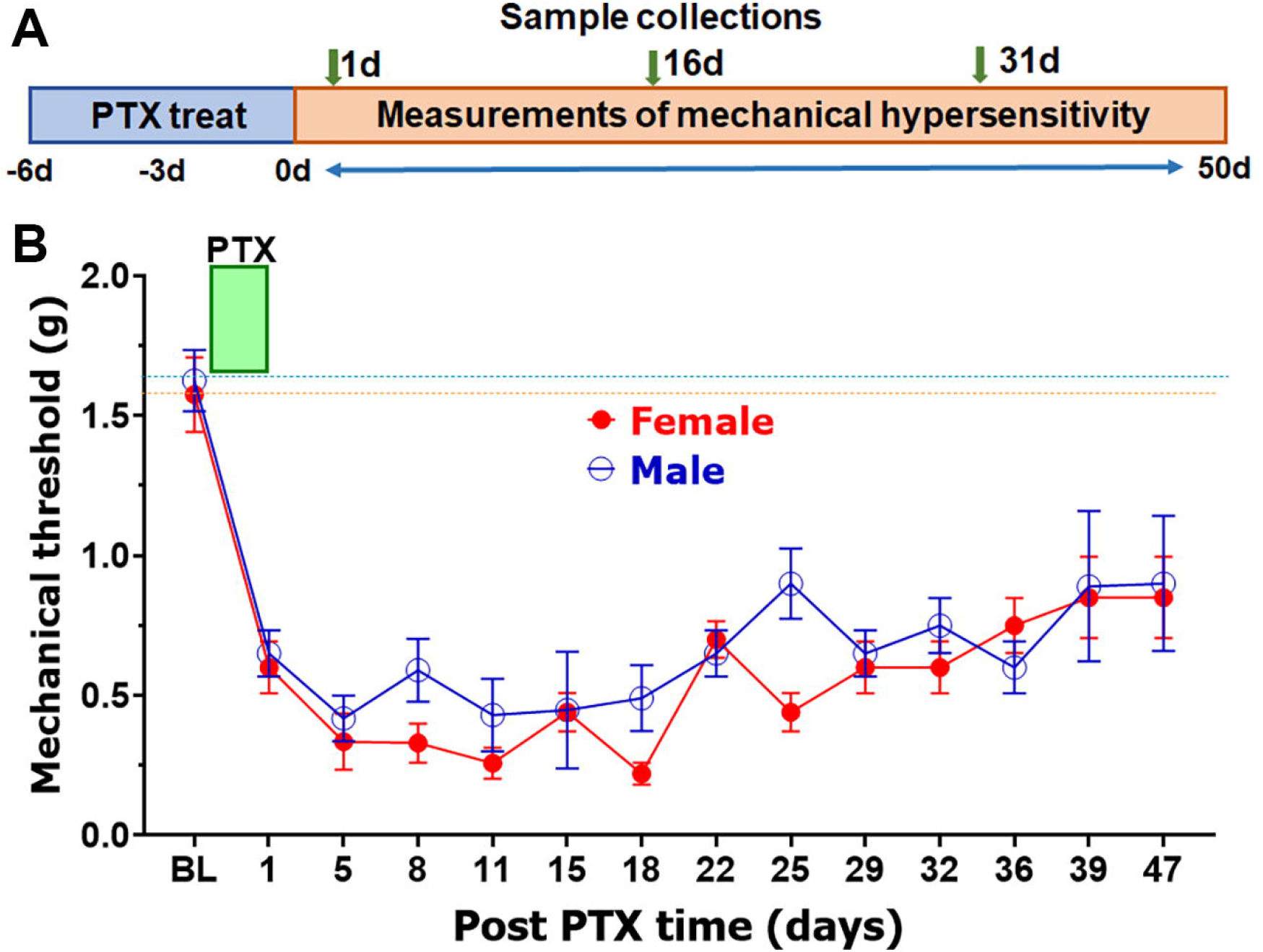
A Paclitaxel (PTX) treatment paradigm for persistent painful chemotherapy-induced pain in males and females. **(A)** Schematic for systemic PTX treatment with 6 mg/kg dosages at -6, -3 and 0 days; and time points for measurements of mechanical hypersensitivity and hind paw and DRG sample collections. **(B)** Time course for the development of mechanical hypersensitivity after systemic PTX treatments of male and female mice.

Mechanical hypersensitivity reflected as the mechanical withdrawal threshold at the hind paw was assessed using the up-down von Frey filament as previously described (Patil et al., 2019). Mice were habituated for 45–60 min prior to measurements. Baseline (BL) readings were taken on the right (ipsi-lateral) hind paw before PTX injection. Hypersensitivity developments were monitored at the ipsilateral hind paws at the post-PTX time points specified in the text and shown in the figures. The behavior experiments were blinded such that the experimenter was not aware of the treatment conditions. We also used randomized designs for behavioral experiments in which animals were randomly assigned to various experimental groups. Additionally, experiments were performed in several trials with small numbers of mice in the groups.

### Biopsy collection and RNA isolation

We used different mouse groups for tissue collection for RNA sequencing and flow cytometry. The development of mechanical hypersensitivity was monitored in mice used for collection of biopsies for RNA sequencing and flow cytometry. To eliminate the contributions of immune cells from blood, all animals for biopsy collections were perfused with cold PBS before tissue dissections. Biopsies for RNA sequencing were collected 4–6 h after the last measurement of mechanical hypersensitivity. Biopsies for flow cytometry were obtained 12–16 h after the last measurement of mechanical hypersensitivity. DRG dissection was performed as described (Patil et al., 2013a; Mecklenburg et al., 2020). Skin biopsies were collected using a 3-mm cylindrical puncher by inserting up to the fat layer deep on the plantar surface of the hind paws (Parisien et al., 2022).

RNA from relatively soft tissues such as DRG was isolated using RNA-easy solution (Qiagen) and a Bead Mill Homogenizer (Omni International, Kennesaw, GA) as previously described (Patil et al., 2013b; Mecklenburg et al., 2020). Isolation of skin RNA required careful slicing/dissing of tissues before the homogenization step, which is detailed in (LoCoco et al., 2020). RNA quality and integrity were checked using an Agilent 2100 Bioanalyer RNA 6000 Nanochip (Agilent Technologies, Santa Clara, CA). RNA quality selected for sequencing had a RIN score of more than 7.

### Bulk RNA sequencing procedures and analysis

Approximately 500 ng of total RNA was used for cDNA library preparation with oligo-dT primers following the Illumina TruSeq preparation guide (Illumina, San Diego, CA) (Mecklenburg et al., 2020). cDNA libraries were quantified, pooled for cBot amplification, and sequenced using a 50-bp single read run with the Illumina HiSeq 3000 platform. The depth of reads was 30-50×10^6^ bp. RNA-seq cDNA libraries were prepared using oligodT according to the SMART-seq-2 protocol (Picelli et al., 2013; Picelli et al., 2014) with previously described modifications (Mecklenburg et al., 2020).

RNA-sequencing data analysis has been presented previously in detail (Mecklenburg et al., 2020). In brief, demultiplexing to generate the FastQ files was done with CASAVA. The RNA-seq data were aligned to the mouse genome build mm9/UCSC hg19 using the TopHat2 default settings. The alignment BAM files were processed using HTSeq-count to obtain the counts per gene in all samples. DESeq-2 was used to identify differentially expressed genes (DEGs) after performing median normalization. Quality control of outliers, intergroup variability, distribution levels, PCA, and hierarchical clustering analysis were performed to statistically validate the experimental data. Multiple test correction was performed using the Benjamini– Hochberg procedure, and an adjusted p-value (Padj) was generated. If not specified in the text selection criteria for DEGs, reads per kilobase of transcript per million mapped reads (RPKM)>1, fold-change (FC)>1.5 and statistically significant DEGs with Padj<0.05. Venn diagrams were generated using https://bioinfogp.cnb.csic.es/tools/venny/. Genes were clustered according to biological processes using PANTHER software - http://www.pantherdb.org/ (Mecklenburg et al., 2020).

### Flow cytometry

Flow cytometry was used to assess the immune cell profiles in tissue biopsies. Single cell suspensions from tissues were created by treating them for 60-90 min (60 min for DRG and 90 min for skin) at 37°C with 250 μg/ml Liberase (Millipore-Sigma; St. Louis, MO) and 100 μg/ml dispase I (Millipore-Sigma; St. Louis, MO), washing with Dulbecco’s modified eagle medium (DMEM) containing 5% fetal calf serum (FCS), triturating with a Pasteur pipette, and then filtering cell suspensions through 70μm strainer.

Cell suspensions were first stained for viability using Zombie NIR™ Fixable Viability Kit (BioLegend; San Diego, CA) for 20 min at room temperature in phosphate buffer solution (pH 7.2S) combined with FcR blocking antibody (1 μg, clone 2.4G2, BioXCell; Lebanon, NH) to block nonspecific binding. Cells were then washed with 2% FBS/PBS and stained with antibodies against surface antigens for 30 min on ice. Fluorochrome-conjugated antibodies against mouse CD45 (clone 30-F11), CD3 (145-2C11), CD24 (M1/69), B220 (RA3-6B2), CD11b (M1/70), CD64 (X54-5/7.1), CD11c (N418), NK1.1 (PK136), TCRβ (H57-597), MHC-II (M5/114.15.2), Ly-6G (1A8), and Ly-6C (KH1.4) were purchased from BioLegend (San Diego, CA), eBioscience (San Diego, CA), or BD Biosciences (San Jose, CA). Flow cytometry was performed using a Celesta or LSRII cytometer (BD Biosciences; San Jose, CA). Data were analyzed using FlowJo LLC v10.6.1 software.

The gating strategy to select immune populations in the skin and DRG was modified from a previously described approach (Yu et al., 2016). Live/singlets/CD45^+^ cells were gated using the markers listed below to define specific cell populations: neutrophils (Nph, CD11b^+^/Ly6G^+^); macrophages (Mph, CD11b^+^/MHCII^hi^/CD64^+^/CD24^lo^/CD11c^-^); inflammatory monocytes (iMo, CD11b^+^/MHCII^lo^/SSC^lo^/ CD64^+^/Ly6C^hi^); CD11b^+^ dendritic cells; (DCs; CD11^b+^/CD64^-^ /CD24^hi^/MHCII^hi^/CD11c^+^); CD11b^-^ DCs (CD11b^-^/CD64^-^/CD24^hi^/MHCII^hi^/CD11c^+^); natural killer cells (NK; NK1.1^+^/TCRβ^-^); B cells (B, B220^+^/ CD11^b-^/CD11^c-^); and T cells (T, CD3^+^/CD11b^-^/CD11c^-^).

### Statistical analysis

Power analysis for RNA-seq experiments on tissue biopsy samples identified that a sample size of 3 for each group of mice attains >80% power for each test to detect at least 1.5-fold significant difference (FC) in gene expression with an estimated standard deviation of 0.6 (considering low mouse to mouse variation) with a false discovery rate (FDR) of 0.05 using a Benjamini–Hochberg test assuming that the actual distribution is normal. Power analysis for behavioral experiments identified that a sample size of 6 for each group of mice achieves >80% power for each test to detect a significant difference.

GraphPad Prism 8.0 (GraphPad, La Jolla, CA) was used for all statistical analyses. Data in the figures are mean ± standard error of the mean (SEM), with “n” referring to the number of animals per group. Differences between groups with two variables were assessed by two-way ANOVA with Bonferroni post hoc tests. A difference was accepted as statistically significant when p<0.05. Interaction F ratios and associated p values are reported.

## Results

### Persistent painful chemotherapy-induced peripheral neuropathy model in males and females

To define biological processes associated with persistent pain in males and females, we first titrated systemic paclitaxel (PTX) dosages to achieve a long-lasting (>50 days) and persistent painful condition in mice (*Fig 1A*). An increase in the cumulative PTX dosage to 18-24 mg/kg led to a gradual increase in chronicity and persistency of mechanical hypersensitivity in male and female hind paws for at least 50 days (*Fig 1B*). A further increase of PTX overall dosage to 40 mg/kg led to an increase in mortality and unpredictable development of pain trajectories (data not shown). Altogether, 3–4 i.p. injections of 6 mg/kg PTX every third day produced long-lasting and persistent mechanical hypersensitivity in both female and male mice (*Fig 1B*).

Chronic pain conditions develop through several phases(Raoof et al., 2018). Presented in *Figure 1A*, the PTX-induced painful neuropathy model could be divided into the following putative stages: an initiation phase from 0–3 days post-PTX; a maintenance phase from 3–25 days post-PTX; and a persistency phase from >25d post-PTX, for males and females (*Fig 1A*). For tissue collection, we selected one time point within each phase. Thus, for the initiation phase, male and female tissues for bulk RNA-seq were collected 1 day after PTX; for the maintenance phase, male and female tissues for bulk RNA-seq were collected 16 days after PTX; and for the persistency phase, male and female tissues for bulk RNA-seq were collected 31 days after PTX.

### Transcriptomic changes in male and female hind paws and DRG 1 day after PTX

Using criteria of RPKM>1, FC>1.5 and Padj<0.05 for determination of DEGs, at 1d post-PTX, 47 such genes were upregulated and 411 downregulated in female DRG, whereas 676 DEGs were upregulated and 852 downregulated in male DRG (*Suppl Figs 1A, 1B*). At the same time point for hind paws, 243 DEGs were upregulated and 469 DEGs were downregulated in females, and 916 DEGs were upregulated and 1447 DEGs were downregulated in males (*Suppl Figs 1C, 1D*). In addition to the substantially higher number of regulated DEGs in males compared with females, PTX treatment led to the regulation of drastically different sets of DEGs in males compared with females for both DRG and hind paw at 1d post-PTX (*Suppl Fig 1*). These differences were so substantial that overlaps in few DEGs were detected (*Suppl Fig 1*). Such sex-dependent regulation levels are in a sharp contrast compared to the naïve condition, where gene expression differences between male and female hind paws and DRG were minimal (25 and 8 DEGs, respectively) (Mecklenburg et al., 2020).

Biological processes at this PTX post-treatment time point were evaluated using the statistical overrepresentation test using PANTHER analysis. Accordingly, the data in *Figures 2* and *3* show that regulated DEGs at 1d post-PTX were almost exclusively linked to increased or decreased developmental processes in a variety of tissues and cell types. Thus, at 1d post-PTX in female DRG, upregulation of developmental processes was marginal (*Fig 2A*), whereas downregulation of development was clear and extensive (*Fig 2B*). These suppressions in developmental processes affected epithelial cells, the vascular system, and neurons, including downregulation of axonogenesis (*Fig 2B*). Downregulation of developmental processes were accompanied by reductions in the expression of genes related to cell migration and communication and extracellular matrix remodeling (*Fig 2B*). In males, PTX treatment led to processes similar to those in female DRG at 1d post-PTX (*Figs 2C, 2D*). However, increases in developmental processes were more robust compared with those in females, and downregulation was not as widespread (*Figs 2C, 2D*).

**Figure 2:**
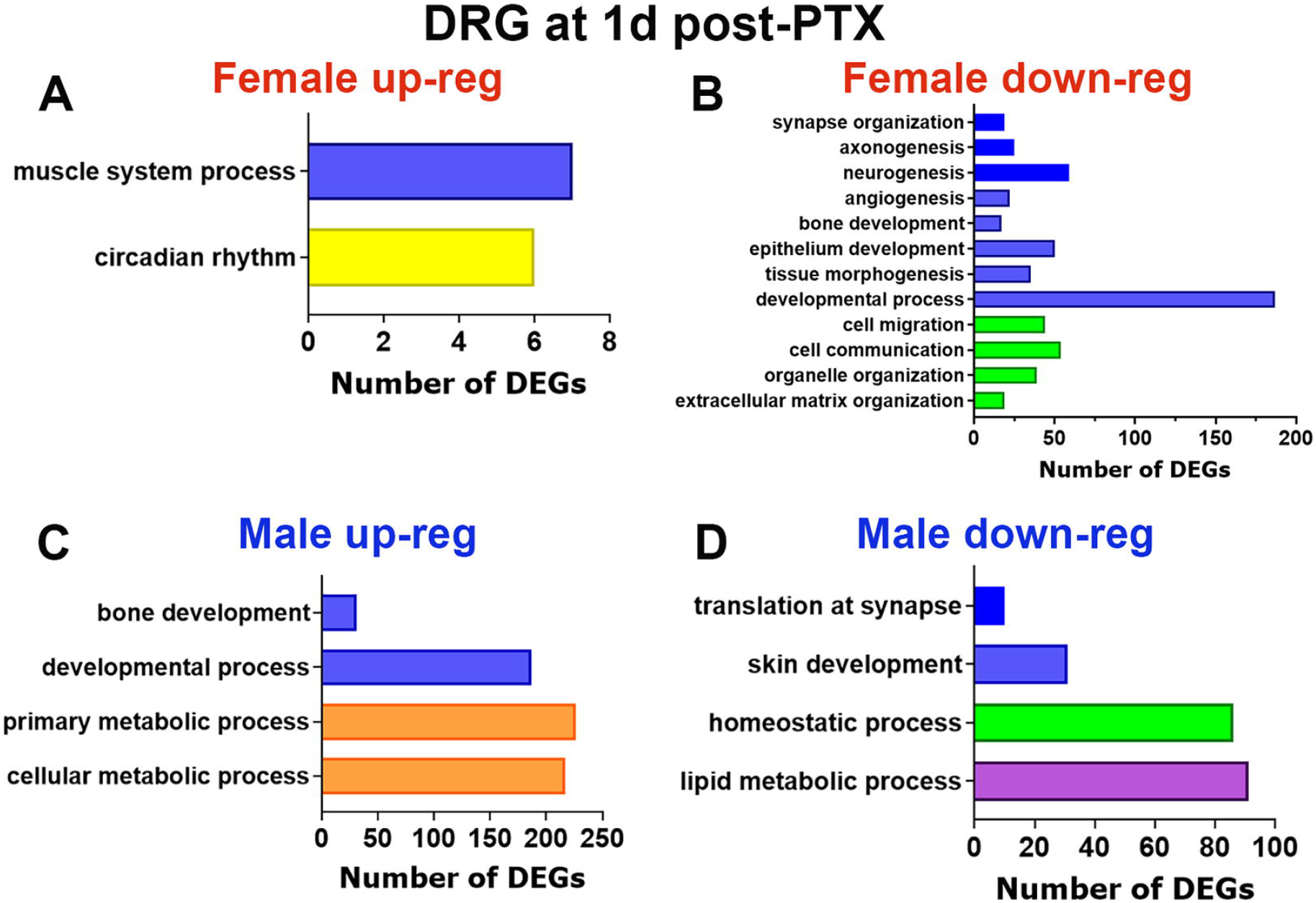
Up- and down-regulated biological processes at 1d post-PTX in female and male DRG. Biological processes for up-regulated (panel **A**) and down-regulated (panel **B**) DEGs in female DRG at 1d post-PTX. Biological processes for up-regulated (panel **C**) and down-regulated (panel **D**) DEGs in male DRG at 1d post-PTX. The X-axis on *the panels A-D* represents numbers of DEGs. The Y-axis notes biological processes.

**Figure 3:**
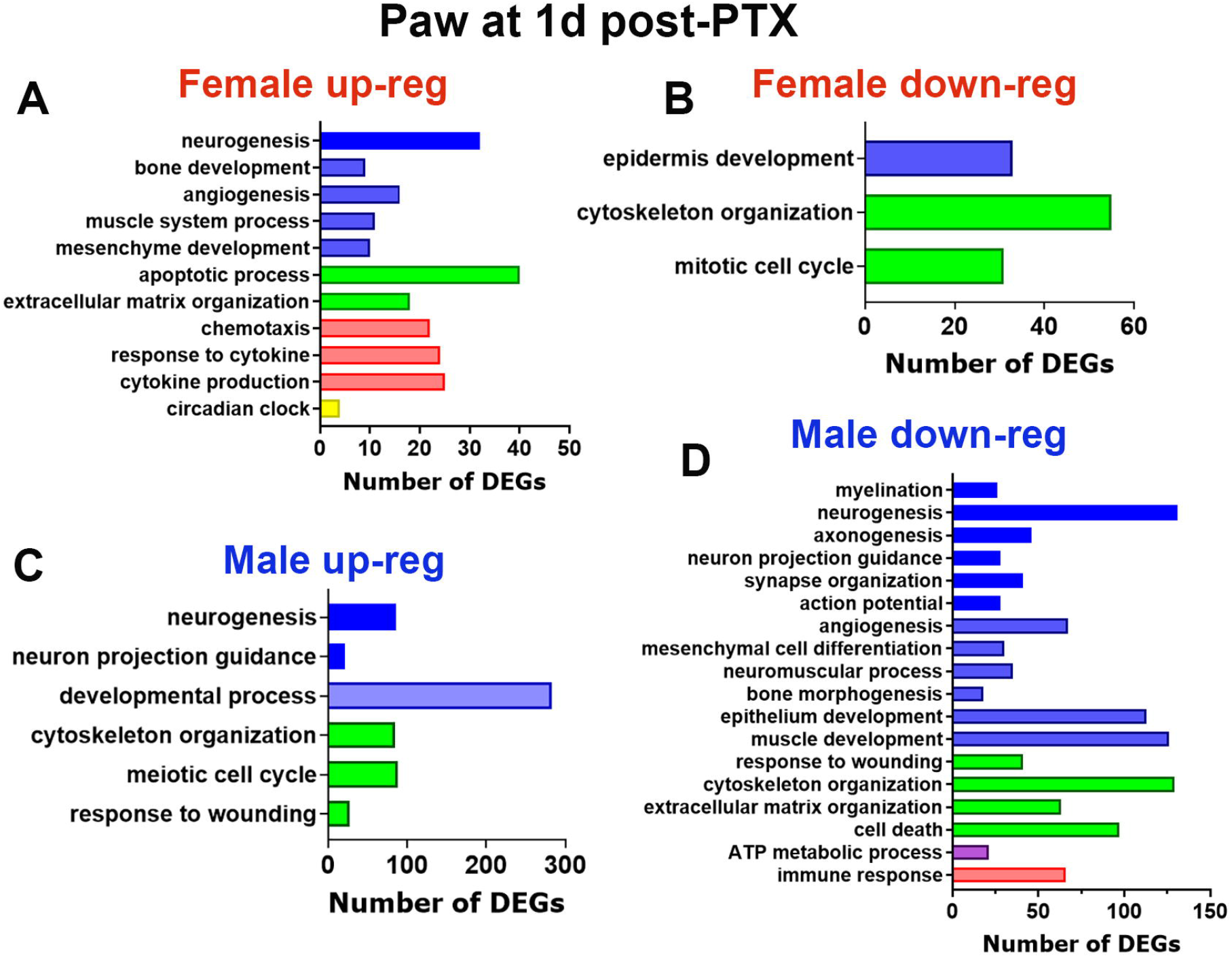
Up- and down-regulated biological processes at 1d post-PTX in female and male hind paws. Biological processes for up-regulated (panel **A**) and down-regulated (panel **B**) DEGs in female hind paws at 1d post-PTX. Biological processes for up-regulated (panel **C**) and down-regulated (panel **D**) DEGs in male hind-paws at 1d post-PTX. The X-axis on *the panels A-D* represents numbers of DEGs. The Y-axis notes biological processes.

In the hind paw, similar processes were detected, and they were again distinct for females versus males (*Fig 3*). Specifically, up-regulation of developmental processes of muscles, the vascular system and neuronal tissues were found, while down-regulation in developmental processes were slight in female hind paws at 1d post-PTX (*Figs 3A, 3B*). In males, the picture was the opposite; downregulation of these processes was found with evidence for genes associated with muscles, epithelium, the vascular system, and neuronal tissues, including a reduction in myelination and axonogenesis (*Figs 3C, 3D*). This down-regulation in developmental processes was accompanied by attenuation in extracellular matrix remodeling, cytoskeleton organization and ATP metabolic process for cellular energy (*Fig 3D*). Interestingly, regulation of DEGs related to the immune system were relatively minimal. Nevertheless, up-regulations in the developmental processes were correlated with an increase of the immune system activity in female paws (*Fig 3A*), while down-regulation of these processes was complemented with a decrease of expression of immune system genes in male paws (*Fig 3D*). Altogether, at the initial stage of post-PTX treatment, regulated DEGs and biological processes were almost exclusively related to up- or downregulation of DEGs related to developmental processes in muscles, epithelium, the vascular system, and neuronal tissues, including myelination, neurogenesis, and axonogenesis. Interestingly, regulation of immune system-related genes was minimal. Furthermore, regulated DEGs, but to a lesser extent biological processes, were substantially different between females and males in paw and DRG tissues.

### Transcriptomic changes at 16 days post-PTX in the hind paws and DRG of males and females

Bulk RNA-seq for hind paw and DRG tissues for males and females showed that at 16 days post-PTX, 51 DEGs were upregulated and 31 downregulated in female DRG, and 108 upregulated and 81 downregulated in hind paws (*Suppl Fig 2*). The numbers of regulated DEGs in males were higher in DRG and similar numbers in paws with 657 up and 388 down in DRG and 147 up and 98 down in paw (*Suppl Fig 2*). Similar to the initiation stage, 16d post-PTX treatment resulted in distinct outcomes in the identities of regulated DEGs for male and female paws and DRG (*Suppl Fig 2*).

Biological processes at the initiation (1 d post-PTX) and maintenance (16d post-PTX) phases were similar in DRG and paws, but there was a reduction in the scope of biological processes and the numbers of regulated DEGs (*Fig 2* vs *Fig 4*; and *Fig 3* vs *Fig 5*). In female DRG, up- and down-regulation of DEGs related to developmental processes became minimal and few biological processes were identified using PANTHER analysis (*Figs 4A, 4B*). In male DRGs, upregulation of the developmental processes for epithelial cells, muscle, and neuronal tissues increased (*Fig 4C*), and downregulation of DEGs representing developmental processes remained at a similar level (*Fig 4D*). In female paws, upregulation of developmental processes was drastically reduced (*Fig 3A* vs *Fig 5A*), and downregulation remained at approximately the same levels (*Fig 3B* vs *Fig 5B*). In male paws, upregulation of developmental processes was stable (*Fig 3C* vs *Fig 5C*), whereas downregulation was reduced (*Fig 3D* vs *Fig 5D*). In male hind paw reduced neurogenesis, axonogenesis, and myelination were still observed (*Fig 5D*). Overall, at 16d post-PTX treatment, DEGs representing developmental processes of a variety of tissues were regulated, but the numbers of DEGs were substantially reduced compared with those at 1d post-PTX with one notable exception: upregulation of developmental processes in male DRG which was substantially increased (*Fig 2C* vs *Fig 4C*).

**Figure 4:**
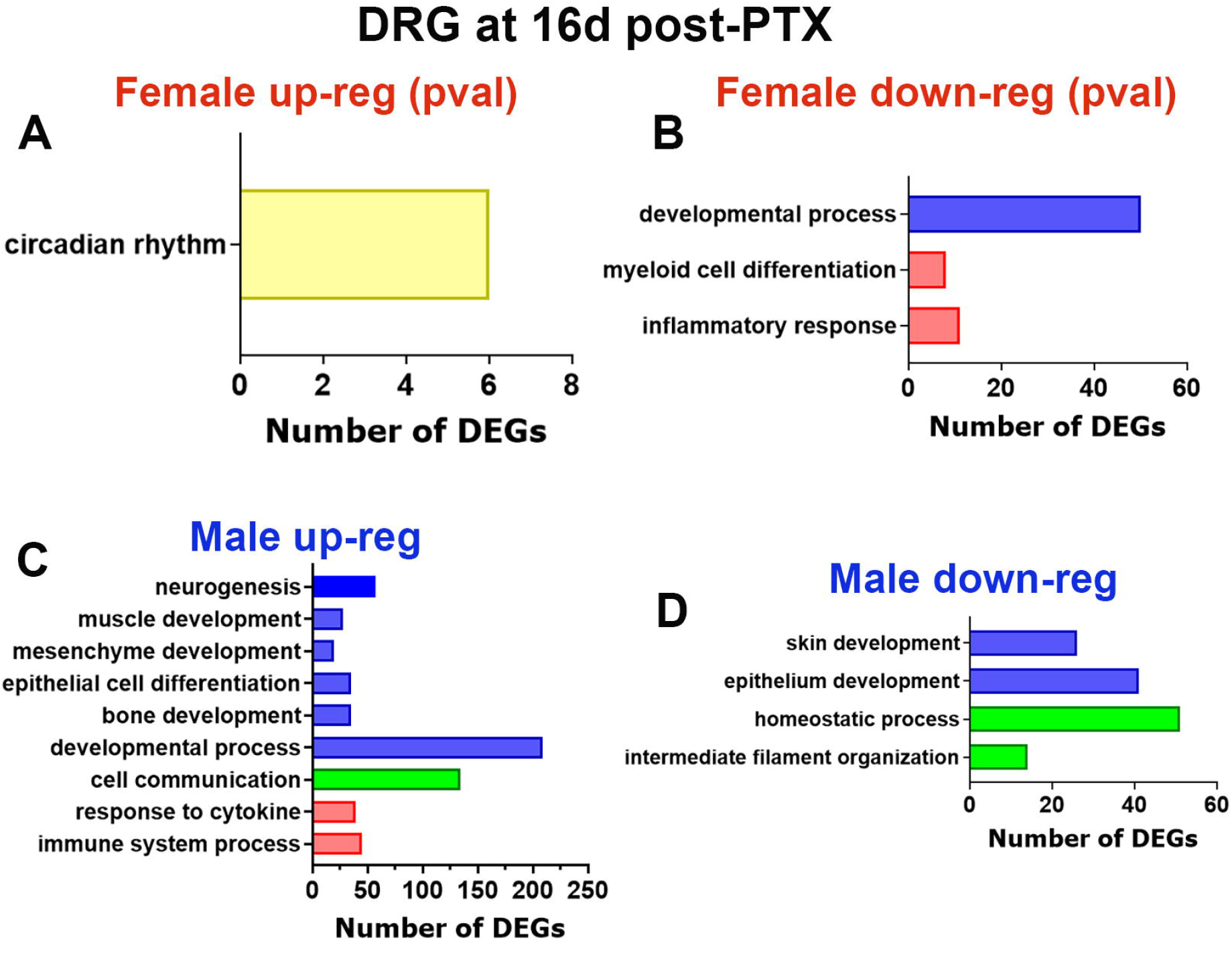
Up- and down-regulated biological processes at 16d post-PTX in female and male DRG. Biological processes for up-regulated (panel **A**) and down-regulated (panel **B**) DEGs in female DRG at 16d post-PTX. Selected DEGs showed statistical difference when Pval<0.05. For other panels in Figures 2-7, selection criteria for DEGs was Padj<0.05. Biological processes for up-regulated (panel **C**) and down-regulated (panel **D**) DEGs in male DRG at 16d post-PTX. The X-axis on *the panels A-D* represents numbers of DEGs. The Y-axis notes biological processes.

**Figure 5:**
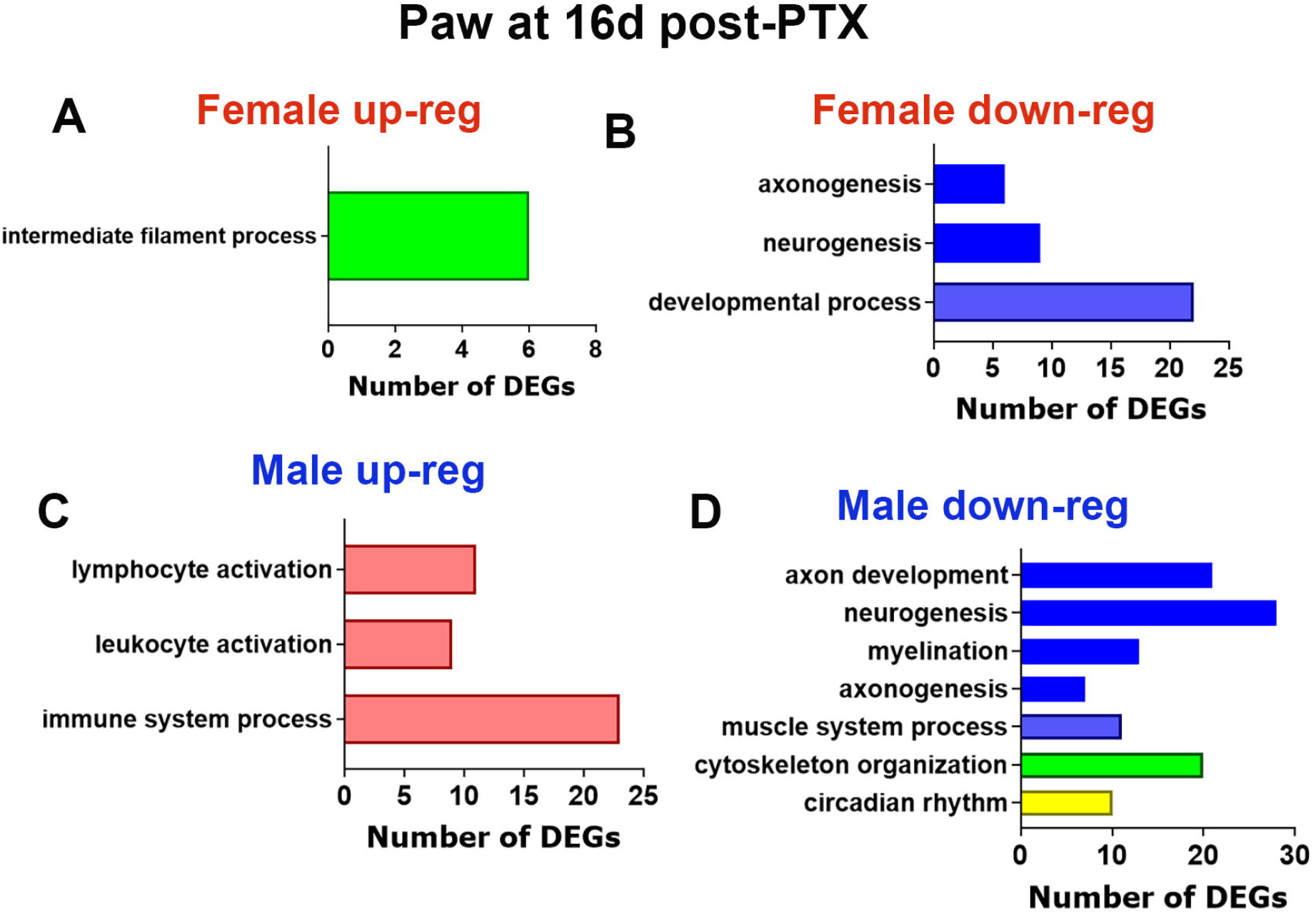
Up- and down-regulated biological processes at 16d post-PTX in female and male hind paws. Biological processes for up-regulated (panel **A**) and down-regulated (panel **B**) DEGs in female hind paws at 16d post-PTX. Biological processes for up-regulated (panel **C**) and down-regulated (panel **D**) DEGs in male hind paws at 16d post-PTX. The X-axis on *the panels A-D* represents numbers of DEGs. The Y-axis notes biological processes.

### Transcriptomic changes at 31 days post-PTX in the DRG and hind paws of males and females

At the 31 days post PTX time point, 7 DEGs were up- and 137 down-regulated in female DRG (*Suppl Figs 3A, 3B*), and only 31 up- and 22 down-regulated were found in the hind paw (*Suppl Figs 3C, 3D*). Numbers of regulated DEGs in male were far higher with 376 up and 314 down-regulated in DRG (*Suppl Figs 3A, 3B*), and 159 up- and 323 down-regulated in hind paws (*Suppl Figs 3C, 3D*). Overlaps between regulated DEGs in female and male DRG and hind paws were low, similar to at 1d and 16d post-PTX (*Suppl Figs 3*). Up-regulated DEGs in female DRG stayed the same as at 16d post-PTX (*Fig 4A vs Fig 6A*). However, downregulated DEGs increased (*Fig 4B vs Fig 6B*). In contrast, up-and downregulated developmental processes in male DRG were similar at 16 and 31 days post-PTX (*Fig 4C vs Fig 6C;* and *Fig 4D vs Fig 6D*). In the hind paw, processes were similar in 16d versus 31d post-PTX for females (*Fig 5A vs Fig 7A;* and *Fig 5B vs Fig 7B*) as well as males (*Fig 5C vs Fig 7C;* and *Fig 5D vs Fig 7D*). Notably, up-regulation of DEGs related to the immune system were yet again related to an increase in developmental processes, while down regulation of immune processes were associated with attenuation an in developmental processes (*Figs 5B, 5C, 7B*). In summary, at 31d post-PTX treatment, up- and downregulation of DEGs representing developmental processes underwent only slight changes from 16 days to 31 days post-PTX in both female and male DRG as well as hind paws (*Figs 6, 7*).

**Figure 6:**
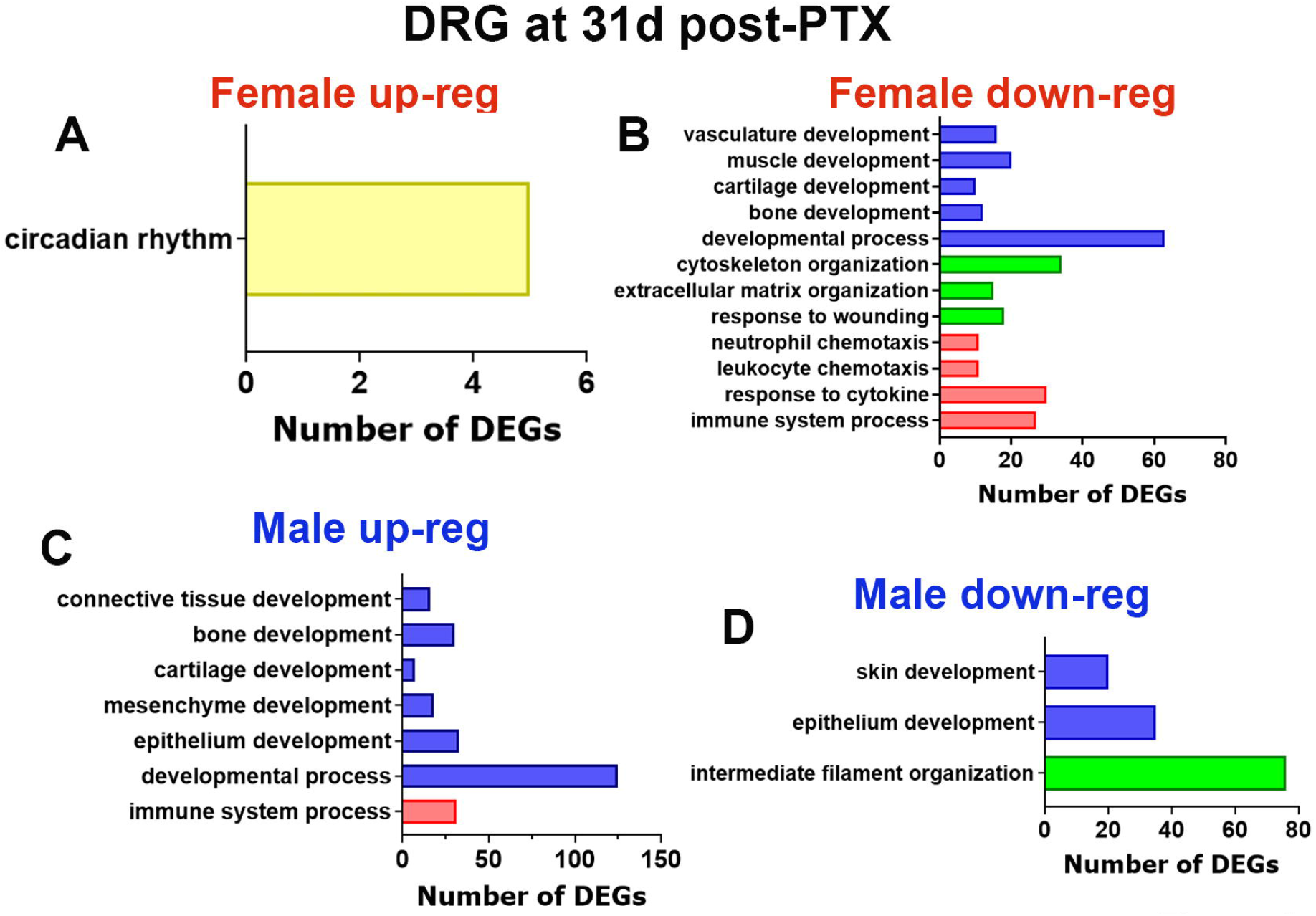
Up- and down-regulated biological processes at 31d post-PTX in female and male DRG. Biological processes for up-regulated (panel **A**) and down-regulated (panel **B**) DEGs in female DRG at 31d post-PTX. Selected DEGs showed statistical difference when Pval<0.05. Biological processes for up-regulated (panel **C**) and down-regulated (panel **D**) DEGs in male DRG at 31d post-PTX. The X-axis on *the panels A-D* represents numbers of DEGs. The Y-axis notes biological processes.

**Figure 7:**
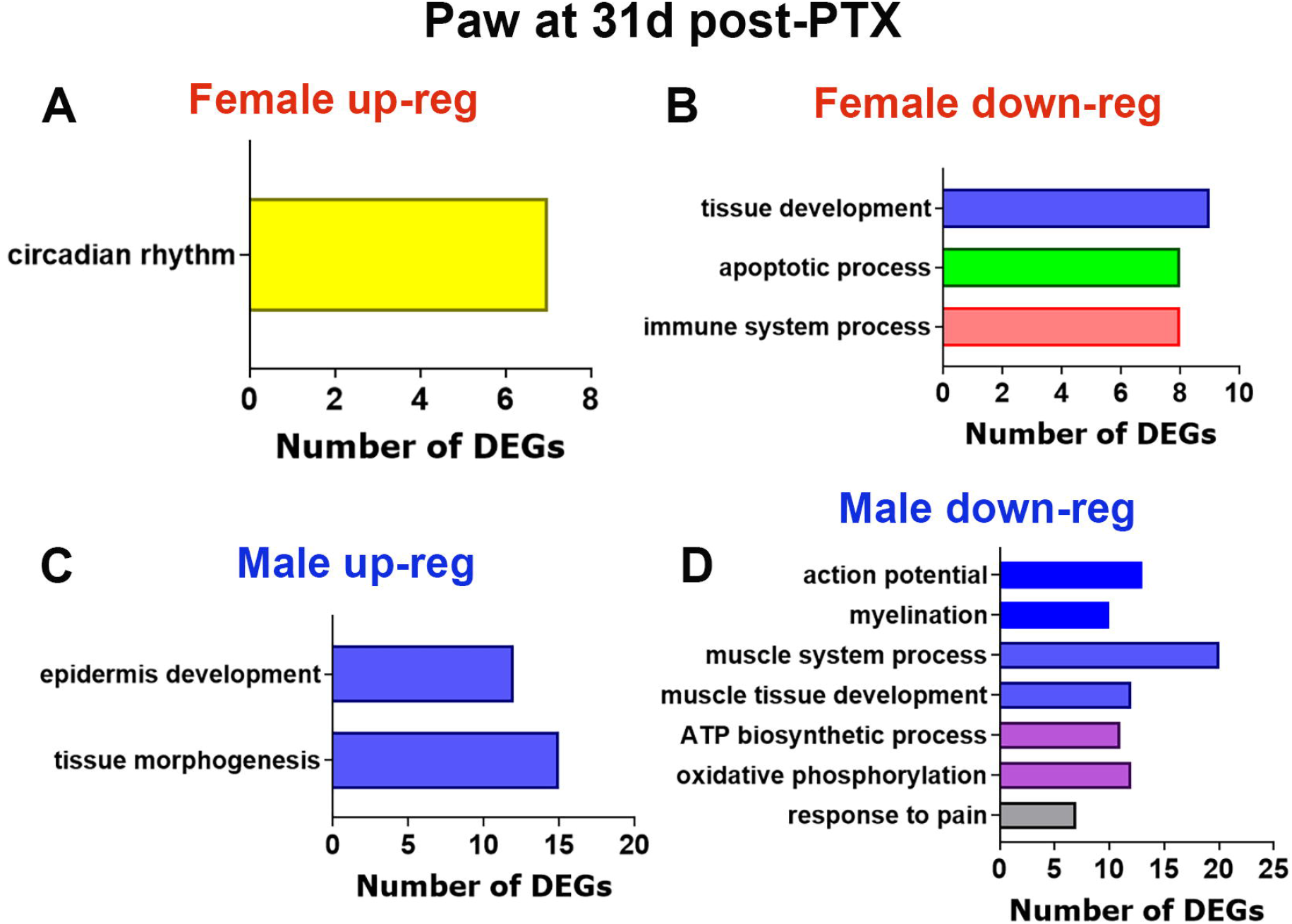
Up- and down-regulated biological processes at 31d post-PTX in female and male hind paws. Biological processes for up-regulated (panel **A**) and down-regulated (panel **B**) DEGs in female hind paws at 31d post-PTX. Selected DEGs showed statistical difference when Pval<0.05. Biological processes for up-regulated (panel **C**) and down-regulated (panel **D**) DEGs in male hind paws at 31d post-PTX. The X-axis on *the panels A-D* represents numbers of DEGs. The Y-axis notes biological processes.

### Interpretation of biological processes during the different phases of persistent neuropathic pain

Gene expression plasticity observed at different phases after PTX treatment was dramatically different for males versus females. Nevertheless, it predominantly represented developmental processes in a variety of nonneuronal and neuronal cell types and/or tissues in both males and females. Somewhat surprisingly, DEG-related proinflammatory processes were lower in number and only slightly changed in both sexes.

Upregulation of DEGs representing the developmental processes of different tissue types could be interpreted as regenerative processes, whereas downregulation of DEGs associated with tissue development could represent degenerative processes. An increase in the expression of genes representing cytoskeleton organization, extracellular matrix remodeling, and oxidative phosphorylation could contribute to tissue and nerve regeneration, whereas a reduction in these processes could reflect degeneration. Thus, muscle and vascular degeneration were associated with the downregulation of DEGs related to muscle development, mitochondrial function, and cytoskeleton organization(Davies et al., 2022; Pan et al., 2022; Duffy et al., 2023; Gallardo et al., 2023). Another example, bulk RNA sequencing revealed that amyotrophic lateral sclerosis (ALS) as a neurodegenerative disease is associated with a reduction in myelination and neurogenesis, synapse organization and transmission, and mitochondrial function(Liu et al., 2020). In contrast, regenerative processes in tissues and nerves were linked to extracellular matrix formation and remodeling, increased axonogenesis, improved tissue differentiation, and increased mitochondrial function (Liedtke et al., 2007; Gao et al., 2020; Luo et al., 2021a; Ronzoni et al., 2021; Wu et al., 2021).

Based on the outlined interpretation framework, we schematically presented the time courses of regenerative and degenerative processes in the hind paw and DRG of males and females (*Fig 8*). Levels of regeneration and degeneration in nonneuronal and neuronal tissues were estimated according to the number of regulated DEGs and the number of biological processes representing degeneration and regeneration. Regeneration and degeneration underwent moderate changes between days 1 and 31 post-PTX (*Fig 8*). The schematics in *Figures 8A* and *8B* also show that the time course of regenerative and degenerative processes distinctly depend on tissue type (DRG vs paw) and sex. Thus, in DRG, degenerative processes persisted through 31d post-PTX in female, and initially affected both nonneuronal (NN) and neuronal (N) tissues (*Fig 8A*). In contrast, in male DRG, regenerative processes dominated, but no balance between these processes was achieved even after 31d post-PTX treatment (*Fig 8A*). In the hind paw, processes were also sex-dependent. Thus, promising regenerative processes at 1d post-PTX were switched to degenerative processes at 16 and 31 days post-PTX treatments in females (*Fig 8B*). In males, a high level of degenerative-associated gene expression changes continued throughout 31d post-PTX and affected both nonneuronal and neuronal tissues (*Fig 8B*).

**Figure 8:**
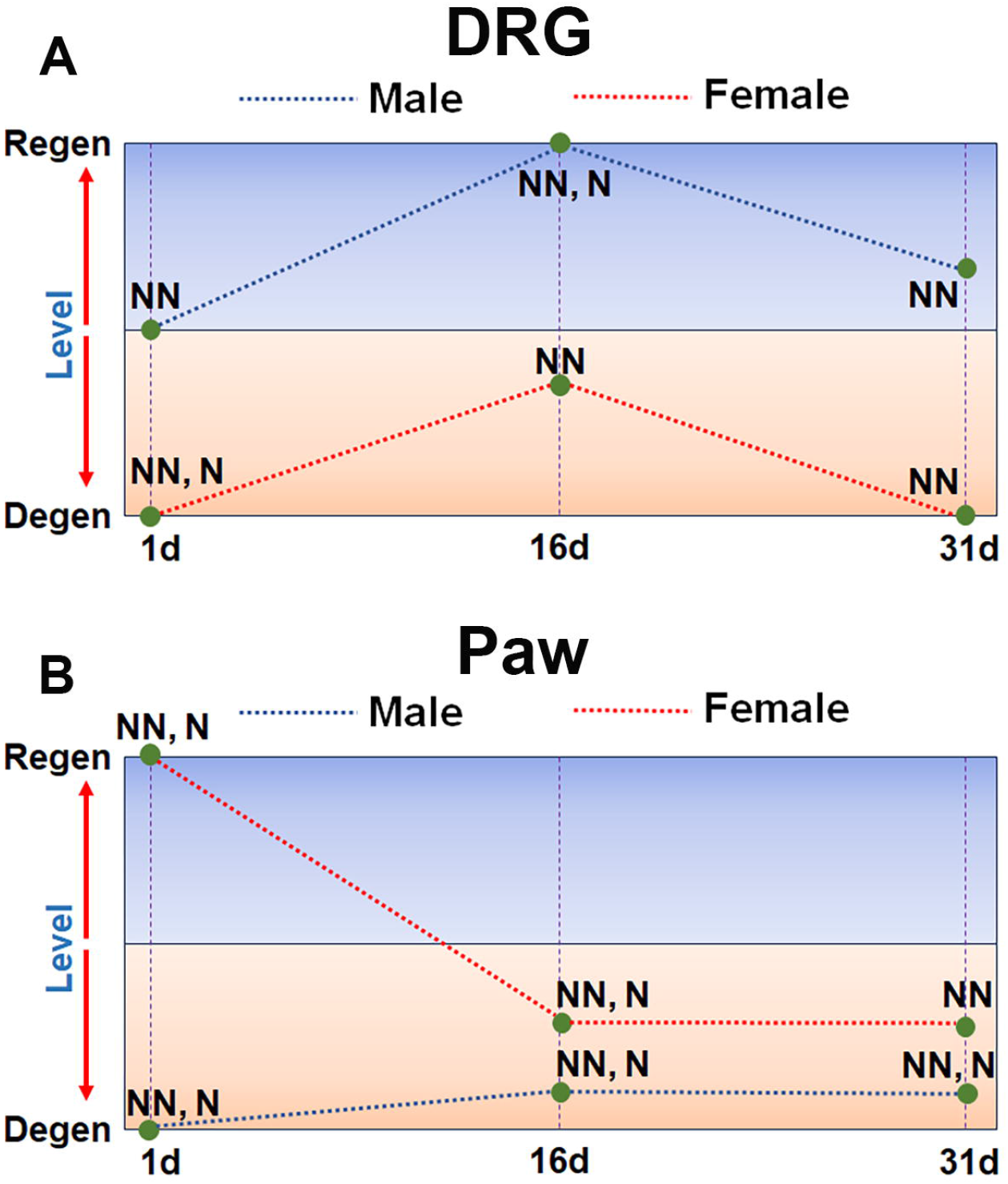
Post-PTX time courses for changes in degenerative and regenerative processes in DRG and hind paw of males and females. **(A)** Schematic representation of degenerative and regenerative processes in DRG of males and females at 1d, 16d and 31d post-PTX. **(B)** Schematic representation of degenerative and regenerative processes in hind paws of males and females at 1d, 16d and 31d post-PTX. The X-axis on *the panels A and B* represents post-PTX days. The Y-axis notes putative levels of degenerative (blue box) or regenerative (orange box) processes. NN – abbreviation for degenerative and regenerative processes in non-neuronal tissues. N – abbreviation for degenerative and regenerative processes in neuronal tissues.

### PTX-induced immune cell profile changes in the male DRG and hind paw

In males, immune system DEGs were upregulated in the DRG at 16d and 31d (*Figs 4C, 6C*) and hind paw at 16d post-PTX (*Fig 5C*); while immune system DEGs were downregulated at 1 day post-PTX in the hind paw (*Fig 3D*). In females, the immune system DEGs were mainly downregulated in the DRG at 16 and 31d (*Figs 4B, 6B*) and hind paw at 31d post-PTX (*Fig 7B*); the exception was upregulation at 1d post-PTX in the hind paw (*Fig 3A*). Detected changes in numbers and expressions of immune system related DEGs were moderate in DRG and hind paws of both males and females. Changes in immune system-related gene expression may be due to immune cell infiltration/ proliferation in tissues and/or activation of resident immune (or potentially non-immune) cells in these tissues.

We investigated whether the regulation of immune-related genes correlated with changes in immune cell profiles in the DRG and hind paw of male and female mice. Flow cytometry was used to examine immune cell profiles in male and female hind paw skin and DRG tissues at different time points after PTX. All immune cell counts were normalized to the number of live “singlet” cells. Overall changes in immune cell (CD45^+^ cells) counts after PTX-treatments were minimal and statistically insignificant in hind paw and DRG of males (1-way ANOVA for paws; F (3, 12) = 0.9914; P=0.4298; n=3-4; 1-way ANOVA for DRG; F (3, 12) = 4.798; P=0.0202; n=3-4; *Suppl Figs 4A, 4B*) and females (1-way ANOVA for paws; F (3, 12) = 2.118; P=0.1512; n=3-4; 1-way ANOVA for DRG; F (3, 12) = 1.883; P=0.1862; n=3-4; *Suppl Figs 4C, 4D*).

Changes in individual immune cell type profiles are presented as normalized total counts (*Figs 9A, 9B, 10A, 10B*) and as counts among CD45^+^ cells (*Figs 9C, 9D, 10C, 10D*). In accordance with previous single-cell RNA-sequencing data, DRG of naïve males and females have had substantial numbers of macrophages (Mph) and neutrophils (Nph) (*Figs 9A, 9C, 10A, 10C*) (Renthal et al., 2020; Mecklenburg et al., 2023). PTX treatment slightly altered immune cell profiles of Mph and Nph, but not inflammatory monocytes (iMo), dendritic (DCs), natural killer (NK), T- and B-cells in DRG of males (2-way ANOVA variables are cell types and post-PTX time points; F (18, 84) = 9.760, p<0.001; n=3-4; *Figs 9A, 9C*). Mph was downregulated at 16d post-PTX and slightly upregulated at 31 days post-PTX (*Fig 9C*), whereas Nph was upregulated only at 31d post-PTX (*Figs 9A, 9C*). This may correspond to minor changes of the immune system-related DEG expressions reflected in *Figures 4C* and *6C*. PTX treatment did not changes iMo, DCs, NK, T- and B-cells in DRG of females (2-way ANOVA variables are cell types and post-PTX time points; *Figs 10A, 10C*).

**Figure 9:**
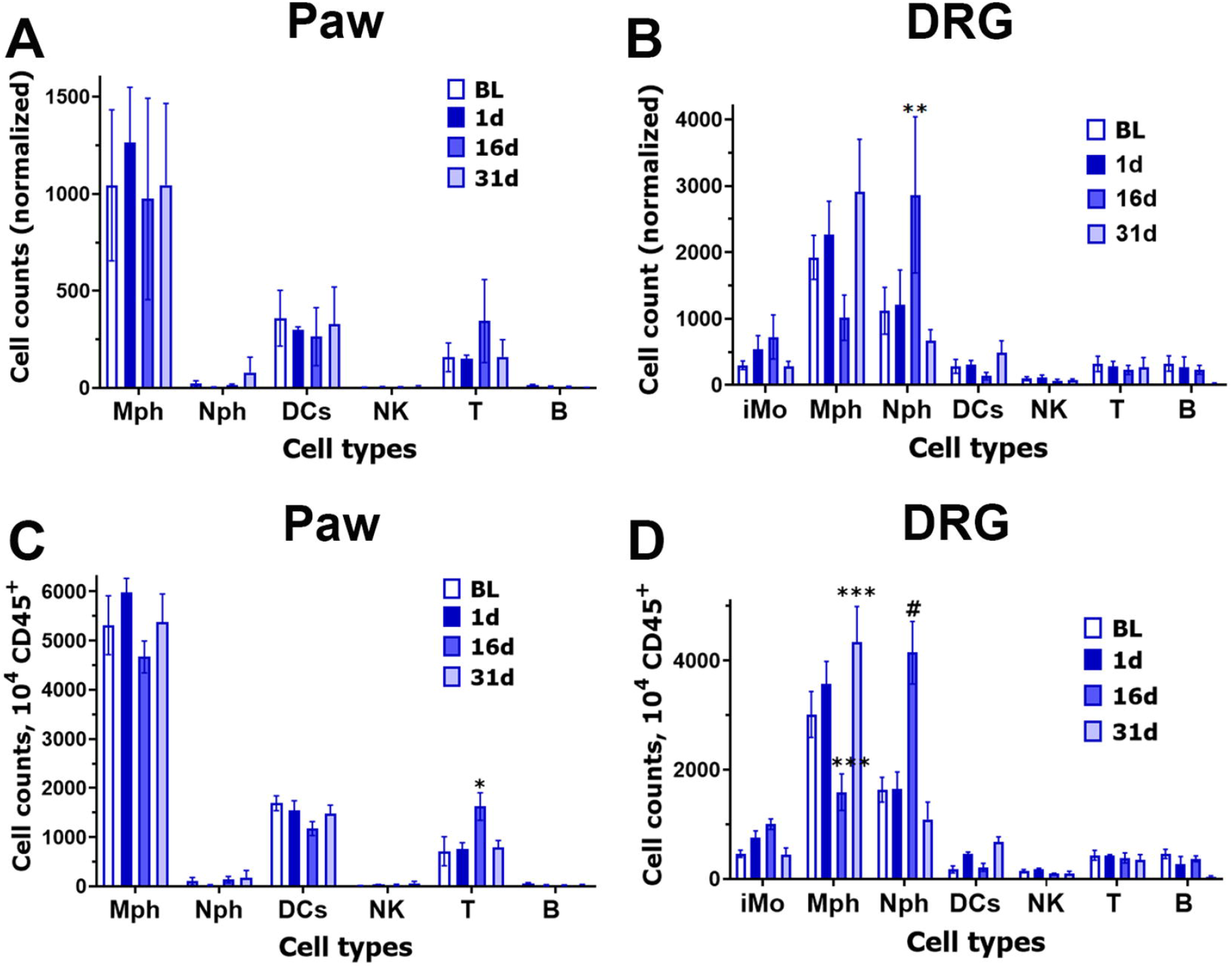
Immune cell profiles in DRG and hind paws of PTX-treated male mice. **(A)** Normalized (by live cells) counts of immune cells in the male DRG at 1, 16 and 31d post-PTX systemic treatments. **(B)** Normalized counts of immune cells in male hind paws at 1, 16 and 31d post-PTX systemic treatments. **(C)** Immune cell counts per 10^4^ CD45^+^ cells in the DRG at 1, 16 and 31d post-PTX systemic treatments. **(D)** Immune cell counts per 10^4^ CD45^+^ cells in the hind paws at 1, 16 and 31d post-PTX systemic treatments. BL is immune cell counts in male DRG and hind paws at 1d post vehicle-treatment. iMo - inflammatory monocytes; Mph – macrophages; Nph – neutrophils; DCs – dendritic cells; NK – natural killer cells; T – T-cells and B – B-cells. Statistic is 2-way ANOVA (* p<0.05; ** p<0.01; *** p<0.001; # p<0.0001; n=3-4).

**Figure 10:**
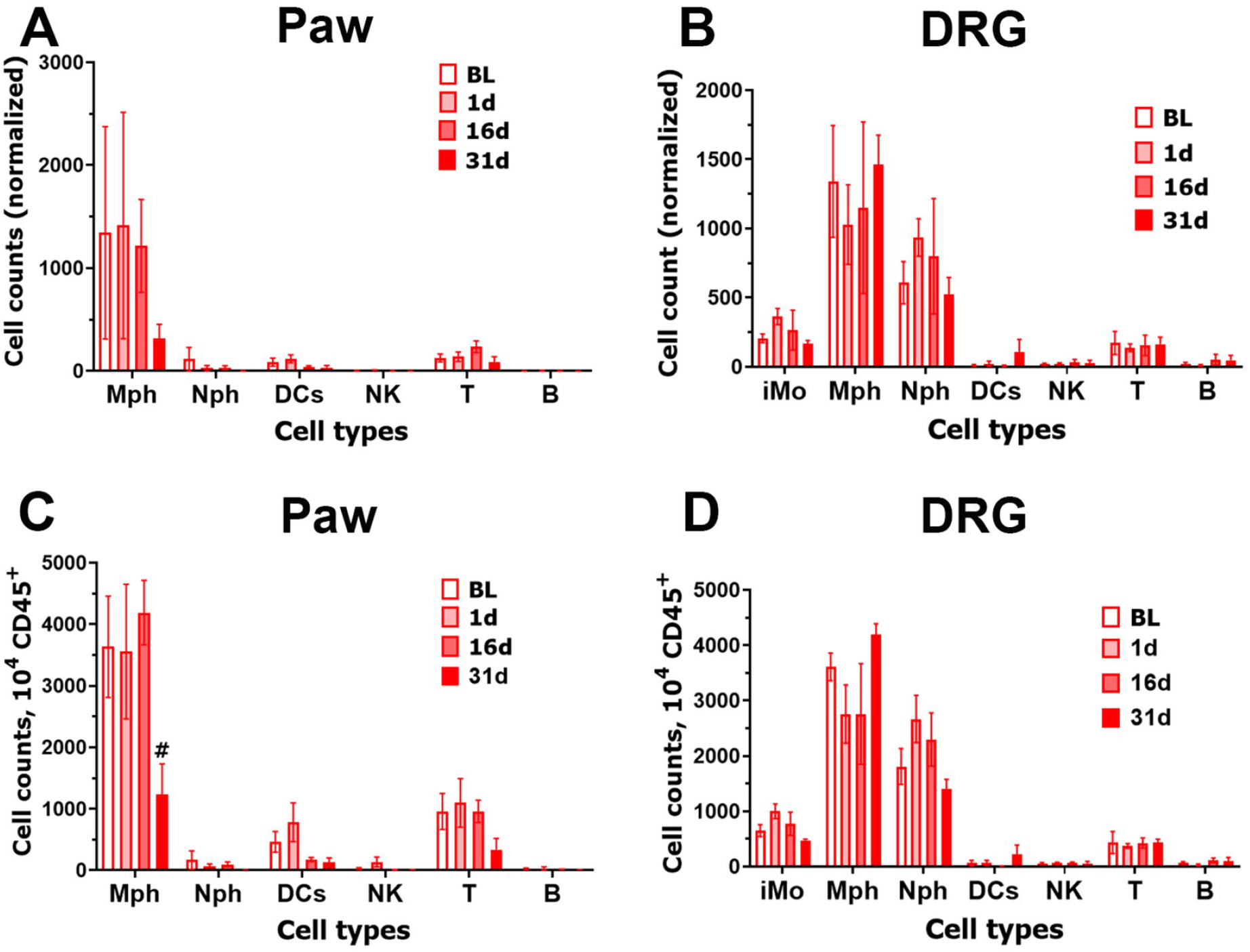
Immune cell profiles in DRG and hind paws of PTX-treated female mice. **(A)** Normalized (by live cells) counts of immune cells in the female DRG at 1, 16 and 31d post-PTX systemic treatments. **(B)** Normalized counts of immune cells in female hind paws at 1, 16 and 31d post-PTX systemic treatments. **(C)** Immune cell counts per 10^4^ CD45^+^ cells in the DRG at 1, 16 and 31d post-PTX systemic treatments. **(D)** Immune cell counts per 10^4^ CD45^+^ cells in the hind paws at 1, 16 and 31d post-PTX systemic treatments. BL is immune cell counts in female DRG and hind paws at 1d post vehicle-treatment. iMo - inflammatory monocytes; Mph – macrophages; Nph – neutrophils; DCs – dendritic cells; NK – natural killer cells; T – T-cells and B – B-cells. Statistic is 2-way ANOVA (# p<0.0001; n=3-4).

Hind paws of naïve males and females had detectable numbers of Mph, DCs, and T cells (*Figs 9B, 9D, 10B, 10D*). PTX treatment altered immune cell profiles only in T cells, but not in Mph, Nph, DCs, NK, and B cells (2-way ANOVA variables are cell types and post-PTX time points; F (15, 72) = 2.017, P=0.025; n=3-4; *Figs 9B, 9D*). T cells were slightly upregulated at 16 days post-PTX (*Fig 9D*). This may correspond to (only 23 DEGs) changes in immune system-related DEG expression, as shown in *Figure 5C*. Immune cell profile did not significantly change in female DRG after PTX-treatment (*Figs 10A, 10C*), despite slight down-regulation of immune-related DEGs in DRG at 16d and 31d post-PTX (*Figs 4B, 6B*). However, Mph cell numbers were reduced in female hind paw (2-way ANOVA variables are cell types and post-PTX time points; F (15, 72) = 2.248; P=0.0118; n=3-4; *Figs 10B, 10D*). Overall, our data indicated that infiltration and/or proliferation of immune cells in male, but not female, DRG and hind paws at different post-PTX time points contributes to the detected slim gene expression plasticity in immune system-related DEGs (*Fig 9*).

## Discussion

Our data indicated that pain persistency after PTX treatment was associated with a variety of continuously ongoing tissue damaging and repairing processes in non-neuronal and neuronal cells in the hind paw and DRG. Overall, individual gene expression plasticity after PTX treatment was substantially sex-dependent. However, in both male and female mice PTX-induced gene expression changes represented tissue damage and repair processes. We did not find evidence that these tissue damage or repair processes were accompanied by substantial inflammatory processes in either the DRG or hind paw of males and female mice treated with this high dose of PTX.

Previous reports have indicated that the immune system could be a major contributor to pain persistency (see review(Ji et al., 2016)). Many reports also implied that the immune system is sex-dependently implicated in each pain development phase, including the chronic pain persistency phase (Fillingim et al., 2009; Melemedjian et al., 2010; Traub and Ji, 2013; Ji et al., 2016; Huh et al., 2017; Price et al., 2018; Raoof et al., 2018; Luo et al., 2021b). For instance, microglia-linked mechanisms lead to neuropathic and inflammatory pain in male rodents but not females(Sorge et al., 2011; Sorge et al., 2015). Activation of an anti-inflammatory IL-13/IL-10 T-cell/macrophage axis in the DRG resolves painful chemotherapy-induced neuropathies (CIPN) in both males and females(Singh et al., 2022). In contrast, stimulation of TLR9 results in CIPN pain initiation only in male mice, whereas an active IL-23/Il-17A/TRPV1 axis promotes pain persistency in females(Luo et al., 2021b). Altogether, reported data indicate that immune cell-mediated mechanisms are sexually dimorphic in chronic pain conditions, leading to sex-dependent promotion of pain chronicity(Hu et al., 2007; Kim et al., 2011; Kobayashi et al., 2015). The general theme emerging from this body of evidence is that males could be more sensitive to the effects of monocyte-lineage cells, whereas females are more sensitive to T cells(Sorge et al., 2015). One commonly held notion is the idea that pain will persist as long as inflammation is present in the DRG and/or spinal cord(Ji et al., 2016). Published data have shown that the elimination of M1 macrophage populations or activation of M2 macrophages in DRG reverses mechanical hypersensitivity in several neuropathic pain models(Mack et al., 2001; Kobayashi et al., 2015; Ji et al., 2016; Zhang et al., 2016; Huh et al., 2017; Laumet et al., 2020; Yu et al., 2020; Luo et al., 2021b). This conclusion is also supported by the fact that most proinflammatory mediators can trigger nociceptive responses in naïve animals by either directly sensitizing sensory neurons or indirectly sensitizing sensory neurons by changing the microenvironment in the vicinity of peripheral and/or central terminals(Ji et al., 2016).

Recently, different mechanisms for pain resolution have been proposed implying that elevation of proinflammatory processes may be required for pain resolution(Parisien et al., 2022). Thus, prolonged intraplantar treatment with prolactin rapidly elevates proinflammatory processes in males, but not in females, and consequently supports pain resolution only in males(Mecklenburg et al., 2022). Elevation of neutrophils in blood and CD11c^+^ microglia in spinal cord resulted in chronic pain resolution in patients and in rodent pain models(Kohno et al., 2022; Parisien et al., 2022). Our data are broadly consistent with this mechanism and show that associations between proinflammatory processes and pain persistency in both males and females are minimal. Despite mechanical hypersensitivity from day 1 onward, macrophage numbers in DRG were not different from baseline (day 2), significantly reduced from baseline (day 16), or moderately increased from baseline (day 35) (*Figs 9 A-D*) and transcriptomic signatures did not reveal major shifts in genes associated with macrophages. We do not rule out the possibility that macrophage populations may be playing different roles at different time points; however, this inconsistent correlation with pain behavior suggests that these macrophage populations are not the only drivers of pain behavior. In this study, pain persistency in male and female mice treated with PTX correlated with either degenerative or regenerative processes in the DRG and hind paw skin. Remarkably, degeneration was always accompanied by a reduction in immune processes, and regeneration was complemented by an increase in immune system activities. This observation could suggest alternative hypothesis that proinflammatory macrophages are reacting to global changes in tissue physiology after PTX treatment and attempting to drive degenerative and regenerative processes toward tissue homeostasis with any/minimal role in driving pain behaviors.

In conclusion, despite PTX-induced persistent painful neuropathy triggering dramatic sex-dependency in gene expression plasticity, biological processes in females and males at every time point consistently involved either degeneration or regeneration processes. These degenerative and regenerative processes affected non-neuronal (angiogenesis, cartilage, bone, muscle and epithelial cell development) as well as neuronal tissues (myelination, axonogenesis and neurogenesis). As expected, these events were accompanied by cytoskeleton and extracellular matrix remodeling, oxidative phosphorylation, and cell energy production. These results add a new depth to our understanding of the biological processes linked to maintaining chronic pain conditions in males and females. Our data could also suggest the potential role of the immune system in tissue repair machinery in DRG and locally at the site where pain occurs.

## Abbreviations

DRG: dorsal root ganglia
CIPN: chemotherapy-induced peripheral neuropathies
PTX: Paclitaxel
i.p.: intra peritoneal injection
i.pl.: intra-plantar injection
BL: baseline
RNA-seq: bulk RNA sequencing
FC: fold change
DEG: differentially expressing gene
RPKM: reads per kilobase of transcript per million mapped reads
FDR: false discovery rate
Padj: an adjusted p-value

## Ethical approval and informed consent

The reporting in the manuscript follows the recommendations in the ARRIVE guidelines (PLoS Bio 8(6), e1000412, 2010). All experimental protocols were approved by the UTHSCSA IACUC committee. Protocol numbers are 20180001AR and 20200011AR.

## Acknowledgements

We would like to thank Mrs Dawn Garcia and Mrs Korri Weldon for their assistance in performing the RNA-seq experiments. The RNA-seq experiments were conducted in the Genome Sequencing Facility (GSF) at the Greehey Children’s Cancer Research Institute (GCCRI) of UTHSCSA. The GSF facility was constructed in part with the support of UT Health San Antonio, NIH/NCI P30 CA054174 (Cancer Center at UT Health San Antonio), NIGMS/NIH S10 Shared Instrumentation Grant Program (SIG) (S10OD021805-01 to Z.L.), and Cancer Prevention Research Institute of Texas (CPRIT) Core Facility Award (RP160732). The Flow Cytometry Shared Resource at UT Health San Antonio is supported by a grant from the National Cancer Institute to the Mays Cancer Center (P30CA054174), a grant from the Cancer Prevention and Research Institute of Texas (CPRIT) (RP210126), a grant from the National Institutes of Health (S10OD030432), and support from the Office of the Vice President for Research at UT Health San Antonio.

## Author Contributions

G.T.N and J.M. conducted the majority of experiments; S.A.S. and Y.Z. performed certain experiments and analyses; G.T.N and J.M. contributed to tissue preparation; G.T.N., Z.L., A.V.T. T.J.P., and A.N.A. analyzed the data, edited the draft, and prepared the final version of the manuscript; A.V.T. T.J.P. and A.N.A. designed and directed the project, wrote the first draft of the manuscript, and prepared the final version of the manuscript. All authors have reviewed the manuscript.

## Data and Material Availability

RNA-seq data have been deposited in GEO Accession, the number GSE240708. Supplementary Excel files show the raw gene readings/counts per gene for all sequencing experiments and the RPKM data for each sample. These supplementary files are “*Veh vs 1d post-PTX (DRG; male)*” for RNA-seq from male mouse DRG at 1d post-PTX; “*Veh vs 16d post-PTX (DRG; male)*” for RNA-seq from male mouse DRG at 16d post-PTX; “*Veh vs 31d post-PTX (DRG; male)*” for RNA-seq from male mouse DRG at 31d post-PTX; “*Veh vs 1d post-PTX (paw; male)*” for RNA-seq from male mouse hind paw at 1d post-PTX; “*Veh vs 16d post-PTX (paw; male)*” for RNA-seq from female mouse DRG at 31d post-PTX; “*Veh vs 31d post-PTX (paw; male)*” for RNA-seq from female mouse DRG at 1d post-PTX; “*Veh vs 1d post-PTX (DRG; female)*” for RNA-seq from female mouse DRG at 1d post-PTX; “*Veh vs 16d post-PTX (DRG; female)*” for RNA-seq from female mouse DRG at 16d post-PTX; “*Veh vs 31d post-PTX (DRG; female)*” for RNA-seq from female mouse DRG at 31d post-PTX; “*Veh vs 1d post-PTX (paw; female)*” for RNA-seq from female mouse DRG at 31d post-PTX; “*Veh vs 16d post-PTX (paw; female)*” for RNA-seq from female mouse DRG at 1d post-PTX; “*Veh vs 31d post-PTX (paw; female)*” for RNA-seq from female mouse hind paw at 31d post-PTX.

