## Supplementary Figures 1-3 for "Degenerative and regenerative peripheral processes are associated with persistent painful chemotherapy-induced neuropathies in males and females"

### Supplementary Figure Legends:

**Supplementary Figure 1: Comparisons of up- and down-regulated DEGs in the female versus male DRG and paw at 1d after PTX treatments**

(**A**) Venn diagram of PTX-induced up-regulation of DEGs in the DRG of females versus males at 1d post-treatments. (**B**) Venn diagram of PTX-induced down-regulation of DEGs in the DRG of females versus males at 1d post-treatments. (**C**) Venn diagram of PTX-induced up-regulation of DEGs in the paws of females versus males at 1d post-treatments. (**D**) Venn diagram of PTX-induced down-regulation of DEGs in the paws of females versus males at 1d post-treatments. Numbers of regulated DEGs in female and male DRG and paws are indicated.

**Supplementary Figure 2: Comparisons of up- and down-regulated DEGs in the female versus male DRG and paw at 16d after PTX treatments**

(**A**) Venn diagram of PTX-induced up-regulation of DEGs in the DRG of females versus males at 16d post-treatments. (**B**) Venn diagram of PTX-induced down-regulation of DEGs in the DRG of females versus males at 16d post-treatments. (**C**) Venn diagram of PTX-induced up-regulation of DEGs in the paws of females versus males at 16d post-treatments. (**D**) Venn diagram of PTX-induced down-regulation of DEGs in the paws of females versus males at 16d post-treatments. Numbers of regulated DEGs in female and male DRG and paws are indicated.

**Supplementary Figure 3: Comparisons of up- and down-regulated DEGs in the female versus male DRG and paw at 31d after PTX treatments**

(**A**) Venn diagram of PTX-induced up-regulation of DEGs in the DRG of females versus males at 31d post-treatments. (**B**) Venn diagram of PTX-induced down-regulation of DEGs in the DRG of females versus males at 31d post-treatments. (**C**) Venn diagram of PTX-induced up-regulation of DEGs in the paws of females versus males at 31d post-treatments. (**D**) Venn diagram of PTX-induced down-regulation of DEGs in the paws of females versus males at 31d post-treatments. Numbers of regulated DEGs in female and male DRG and paws are indicated.

**Supplementary Figure 4: CD45 cell in DRG and hind paws of PTX-treated male and female mice.**

**(A)** Normalized (by live cells) counts of CD45 cells in the male hind paws at 1, 16 and 31d post-PTX systemic treatments. **(B)** Normalized counts of CD45 cells in the male DRG at 1, 16 and 31d post-PTX systemic treatments. **(C)** Normalized counts of CD45 cells in the female hind paws at 1, 16 and 31d post-PTX systemic treatments. **(D)** Normalized counts of CD45 cells in the female DRG at 1, 16 and 31d post-PTX systemic treatments. BL is CD45 cell counts in DRG and hind paws at 1d post vehicle-treatment. Tissue types and sexes are indicated above each panel.

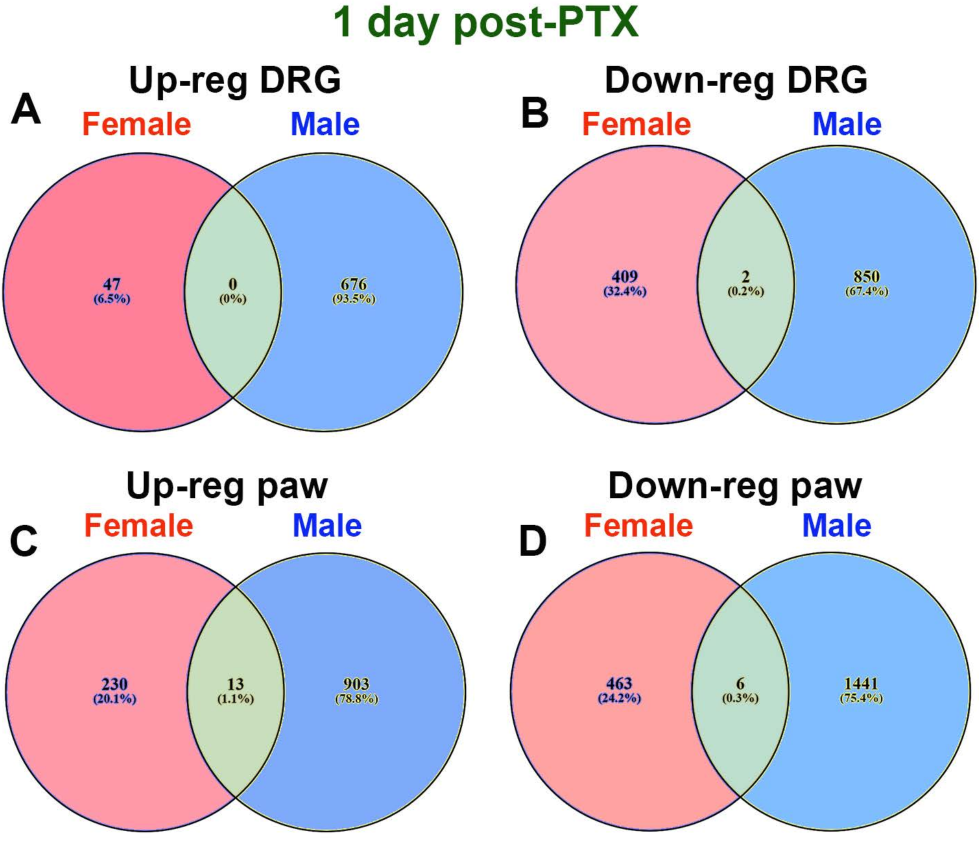
1 day post-PTX
A
Up-reg DRG
Female
Male
47
(6.5%)
0
(0%)
676
(93.5%)
B
Down-reg DRG
Female
Male
409
(32.4%)
2
(0.2%)
850
(67.4%)
C
Up-reg paw
Female
Male
230
(20.1%)
13
(1.1%)
903
(78.8%)
D
Down-reg paw
Female
Male
463
(24.2%)
6
(0.3%)
1441
(75.4%)

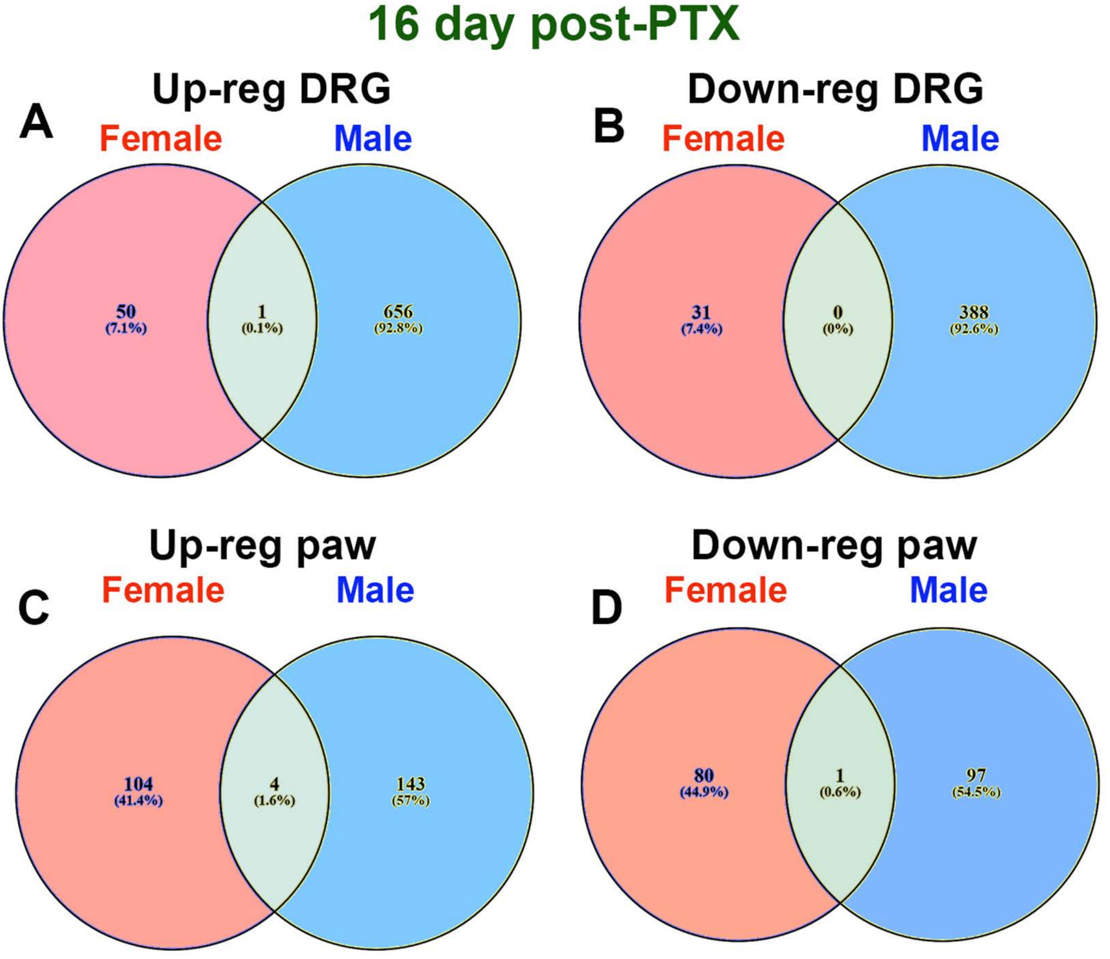
16 day post-PTX
A
Up-reg DRG
Female
Male
50
(7.1%)
1
(0.1%)
656
(92.8%)
B
Down-reg DRG
Female
Male
31
(7.4%)
0
(0%)
388
(92.6%)
C
Up-reg paw
Female
Male
104
(41.4%)
4
(1.6%)
143
(57%)
D
Down-reg paw
Female
Male
80
(44.9%)
1
(0.6%)
97
(54.5%)

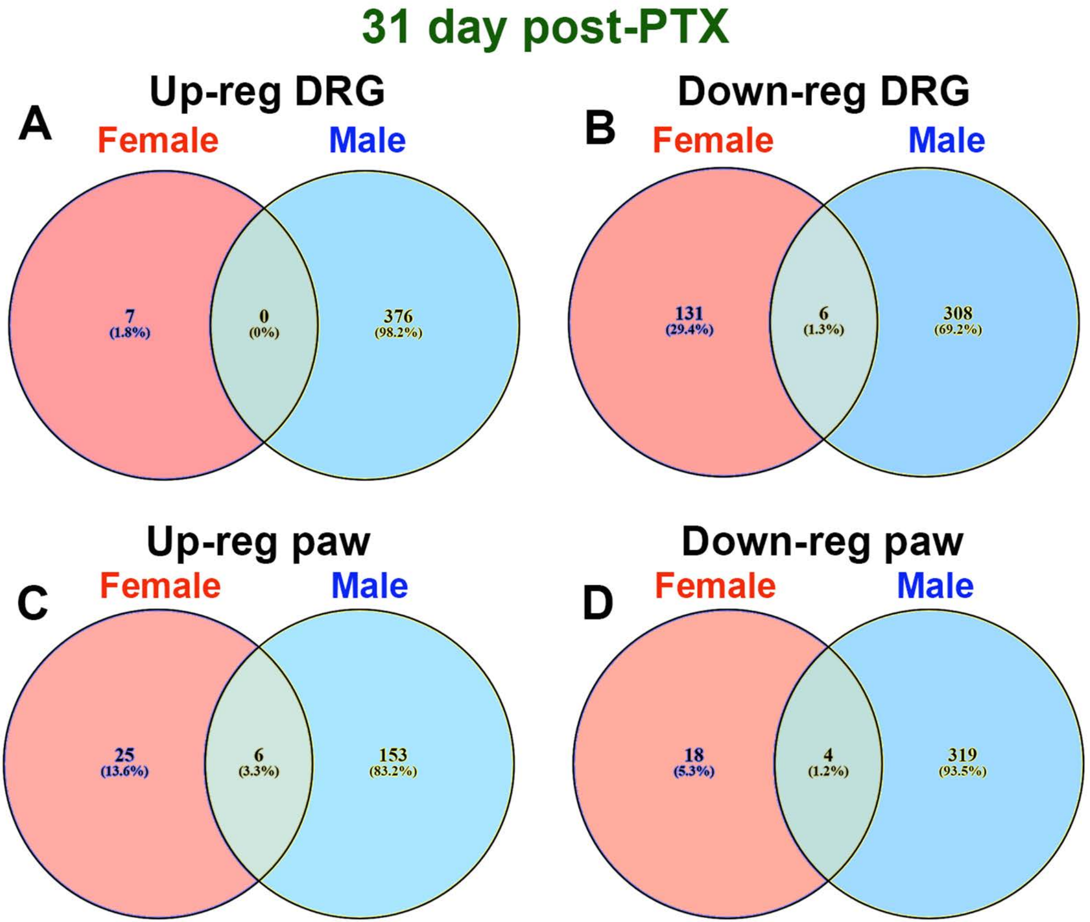

31 day post-PTX
A
Up-reg DRG
Female
Male
7
(1.8%)
0
(0%)
376
(98.2%)
B
Down-reg DRG
Female
Male
131
(29.4%)
6
(1.3%)
308
(69.2%)
C
Up-reg paw
Female
Male
25
(13.6%)
6
(3.3%)
153
(83.2%)
D
Down-reg paw
Female
Male
18
(5.3%)
4
(1.2%)
319
(93.5%)

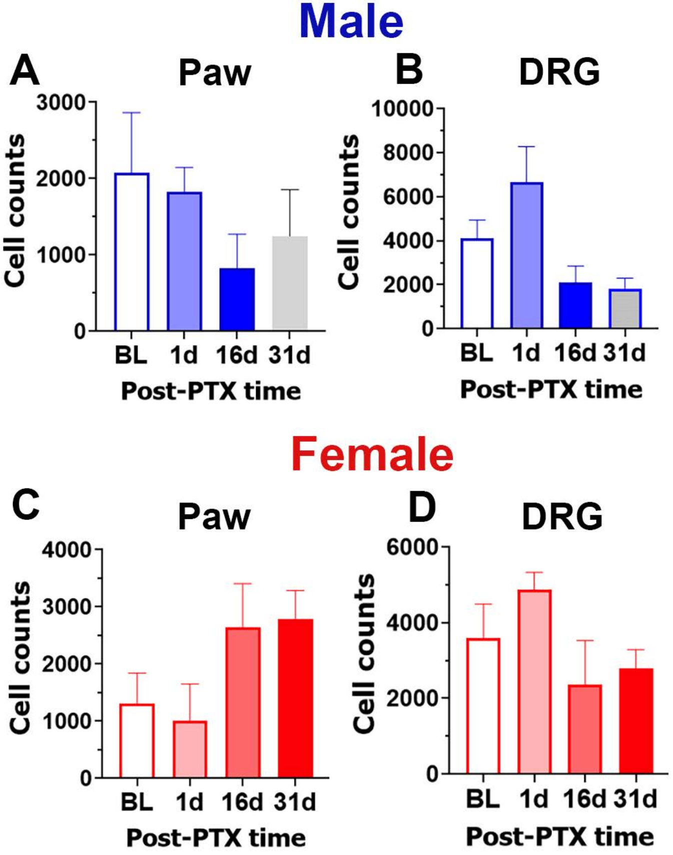

Male
A
Paw
3000
2000
1000
0
Cell counts
BL
1d
16d
31d
Post-PTX time
B
DRG
10000
8000
6000
4000
2000
0
Cell counts
BL
1d
16d
31d
Post-PTX time
Female
C
Paw
4000
3000
2000
1000
0
Cell counts
BL
1d
16d
31d
Post-PTX time
D
DRG
6000
4000
2000
0
Cell counts
BL
1d
16d
31d
Post-PTX time
